# Mutation bias shapes gene evolution in *Arabidopsis thaliana*

**DOI:** 10.1101/2020.06.17.156752

**Authors:** J. Grey Monroe, Thanvi Srikant, Pablo Carbonell-Bejerano, Moises Exposito-Alonso, Mao-Lun Weng, Matthew T. Rutter, Charles B. Fenster, Detlef Weigel

## Abstract

Classical evolutionary theory maintains that mutation rate variation between genes should be random with respect to fitness ^1–4^ and evolutionary optimization of genic mutation rates remains controversial ^3,5^. However, it has now become known that cytogenetic (DNA sequence + epigenomic) features influence local mutation probabilities ^6^, which is predicted by more recent theory to be a prerequisite for beneficial mutation rates between different classes of genes to readily evolve ^7^. To test this possibility, we used de novo mutations in Arabidopsis thaliana to create a high resolution predictive model of mutation rates as a function of cytogenetic features across the genome. As expected, mutation rates are significantly predicted by features such as GC content, histone modifications, and chromatin accessibility. Deeper analyses of predicted mutation rates reveal effects of introns and untranslated exon regions in distancing coding sequences from mutational hotspots at the start and end of transcribed regions in A. thaliana. Finally, predicted coding region mutation rates are significantly lower in genes where mutations are more likely to be deleterious, supported by numerous estimates of evolutionary and functional constraint. These findings contradict neutral expectations that mutation probabilities are independent of fitness consequences. Instead they are consistent with the evolution of lower mutation rates in functionally constrained loci due to cytogenetic features, with important implications for evolutionary biology^8^.

---

A core maxim of evolutionary biology, codified through experiments completed in the early 1940s ^1^, is that the “consequences of a mutation have no influence whatsoever on the probability that this mutation will or will not occur.” ^2^ This assertion has profound implications for understanding organismal evolution and predicting human disease. That genic mutation rates might have been optimized during evolution has been extensively challenged with a strong argument: selection for mutation rates on a gene-by-gene basis cannot overcome the barrier of genetic drift ^3^. And while reports of non-random relationships between mutation rates and fitness consequences have been previously made, these have been questioned because they have largely relied on substitution rates in natural populations rather than direct measures of *de novo* mutations ^3,9–12^.

More recently though, discoveries in genome biology have inspired a reevaluation of classical theories of mutation rate evolution. It is now recognized that mutation rates across genomes are influenced by DNA sequence composition, epigenomic features, and bias in the targets of DNA repair mechanisms ^5,6,13–32^. It is also known that broad classes of genes (e.g., housekeeping genes) exist in distinct cytogenetic (DNA sequence + epigenomic) states. This provides an opportunity for mutation rates to evolve in beneficial directions (e.g., lower mutation rates in cytogenetic states characteristic of essential housekeeping genes where mutations are more likely to be deleterious). Indeed, processes facilitating reduced mutation rates in genic regions ^22,29,33^ and active genes ^15,17,21,24,34^ have already been documented in recent years. These discoveries are consistent with contemporary theoretical predictions from population genetics that beneficial mutation rates could readily evolve even in the face of genetic drift if mutation rates are linked to gene regulation or common sequence and epigenomic features ^3,8^.

Nevertheless, given the historical backdrop of these recent functional discoveries, the possibility of beneficial genic mutation rates (i.e. low rates in functionally constrained genes) remains in conflict with prevailing evolutionary thought. This study aims to synthesize the functional mechanisms and evolutionary implications of mutation bias within and between genes, to ask if such bias shapes patterns of genic evolution and if mutational probabilities are indeed independent of mutational consequences.

The major technical barrier to resolving uncertainty about the evolutionary significance of intra-genomic mutation rate variation has been the limitation of data describing the distribution of new mutations before they have been subject to strong selection in natural populations. We set out to address this challenge using *de novo* mutations and sequence and epigenomic features plausibly linked to mutation rates to create a high resolution predictive model of mutation rate variation across the genome of the model plant *Arabidopsis thaliana.* We first reanalyzed *A. thaliana* mutation accumulation lines ^6^, identifying both putative germline and somatic mutations (Fig. 1a, Extended Data Fig. 1–2, Supplementary Data 1). Strict filtering was used to eliminate false positives – fewer than 10% of all called variants were included in the final high confidence set of *de novo* mutations (Methods). These mutations (n = 10,590, ~20% germline, ~80% somatic, ~69% SNVs, ~31% InDels) are a > 5-fold increase over previous benchmarks ^6,28,35^ to characterize the mutational landscape of the *A. thaliana* genome (Fig. 1b,c).

**Figure 1.**
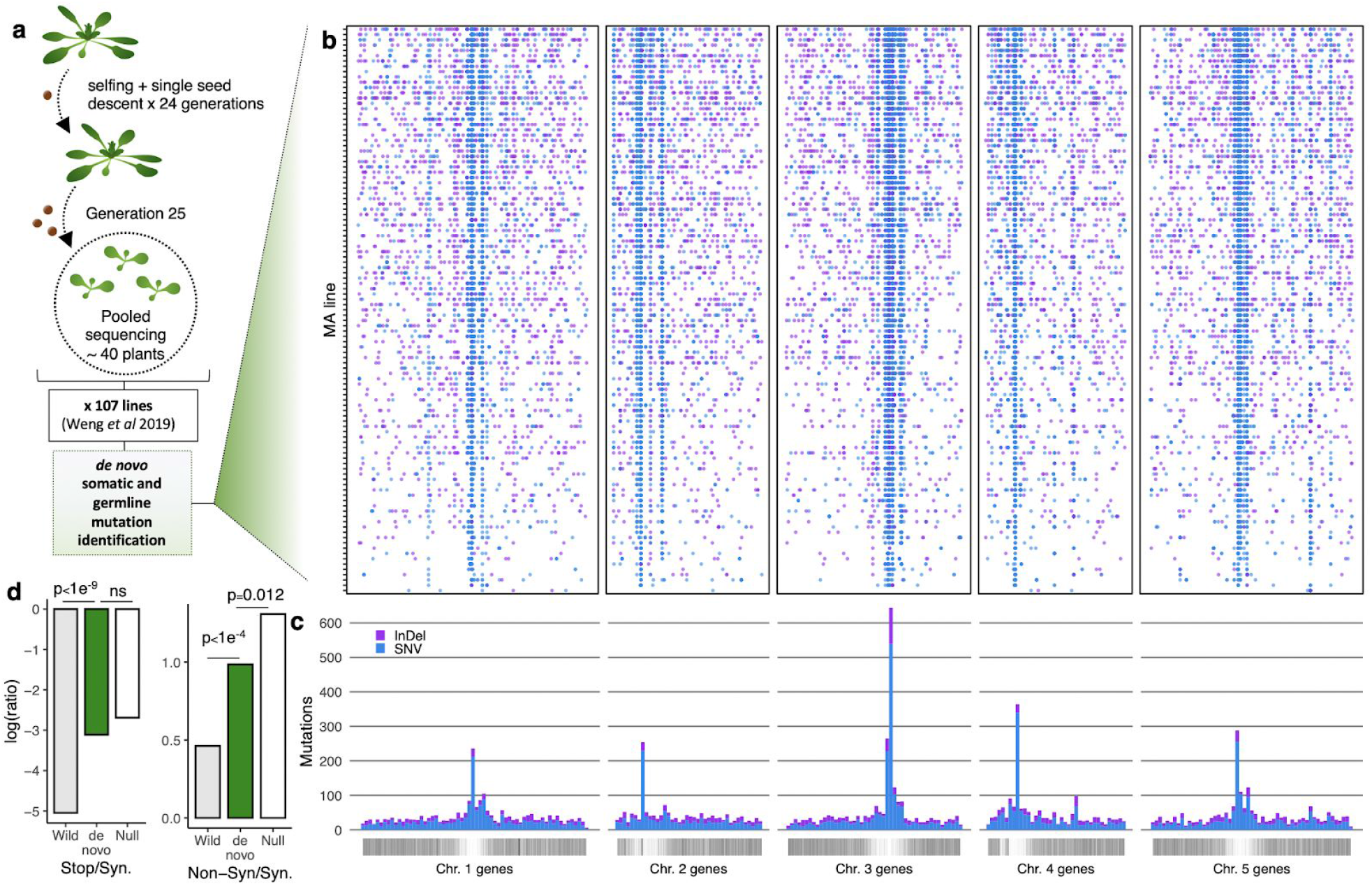
*De novo* mutations across the *Arabidopsis thaliana* genome. (a) Experimental design of mutation data analyzed here. 107 mutation accumulation lines were propagated by single-seed descent for 24 generations before pooled sequencing of seedlings from the 25th generation 6. (b) Locations of mutations detected across 107 lines. Blue points mark the location of single nucleotide variants (SNVs), InDels are marked with purple. (c) Genomic regions of low gene density are enriched in mutations, similar to levels of natural polymorphism (Extended Data Fig. 4). Frequency distribution of *de novo* mutations across the *A. thaliana* nuclear genome. Vertical black lines below mark the location of genes. (d) The *de novo* mutations detected show evidence of being subject to relaxed purifying selection, with elevated rates of non-synonymous and stop codon variants compared to polymorphisms detected in 1,135 natural populations of *A. thaliana*, but lower rates than a null model based on mutation spectra and nucleotide composition of *A. thaliana* coding sequences.

The germline mutations studied here were accumulated with population bottlenecks to N_e_ = 1 at each generation, so only mutations causing lethality or sterility are expected to be removed by selection ^6^. Previous work has shown that somatic mutations in plants experience minimal selection ^36,37^. Thus as expected, the *de novo* mutations identified exhibit significantly relaxed purifying selection with rates of non-synonymous and premature stop codon variants greater than observations in natural populations of *A. thaliana* and closer to a null model (Fig. 1d). Still, to confirm that selection during mutation accumulation in coding regions has not influenced conclusions here, we repeated analyses while explicitly ignoring coding regions when building genic level predictive models and found that results were qualitatively unchanged (Extended Data Fig. 3). We also repeated analyses with only putative germline and somatic mutations, and consistent with plants lacking a completely segregated germline^38^, the general conclusions in relation to the genic distribution of *de novo* mutations proved robust to this subsampling (Extended Data Fig. 3).

This catalog of *de novo* mutations provided data to build predictive models of mutation rates across the *A. thaliana* genome coupled with DNA sequence and epigenetic features. To do so we compiled additional genome wide data characterizing GC content, cytosine methylation, histone modifications, and chromatin accessibility as previous work indicates that such features are probable causes of intra-genomic mutation rate variation. We found that these features could jointly explain > 65% of the variance in regional (100 kb) mutation rates in *A. thaliana* (Extended Data Fig. 4). However, the ultimate aim of this investigation was to study mutation rates at gene level resolution. We therefore calculated the values of these predictive features in all genic regions – all upstream (1 kb), 5’ untranslated regions (UTRs), coding regions, introns, 3’ UTRs, and downstream (1 kb) regions throughout the genome (Fig. 2a,b).

**Figure 2.**
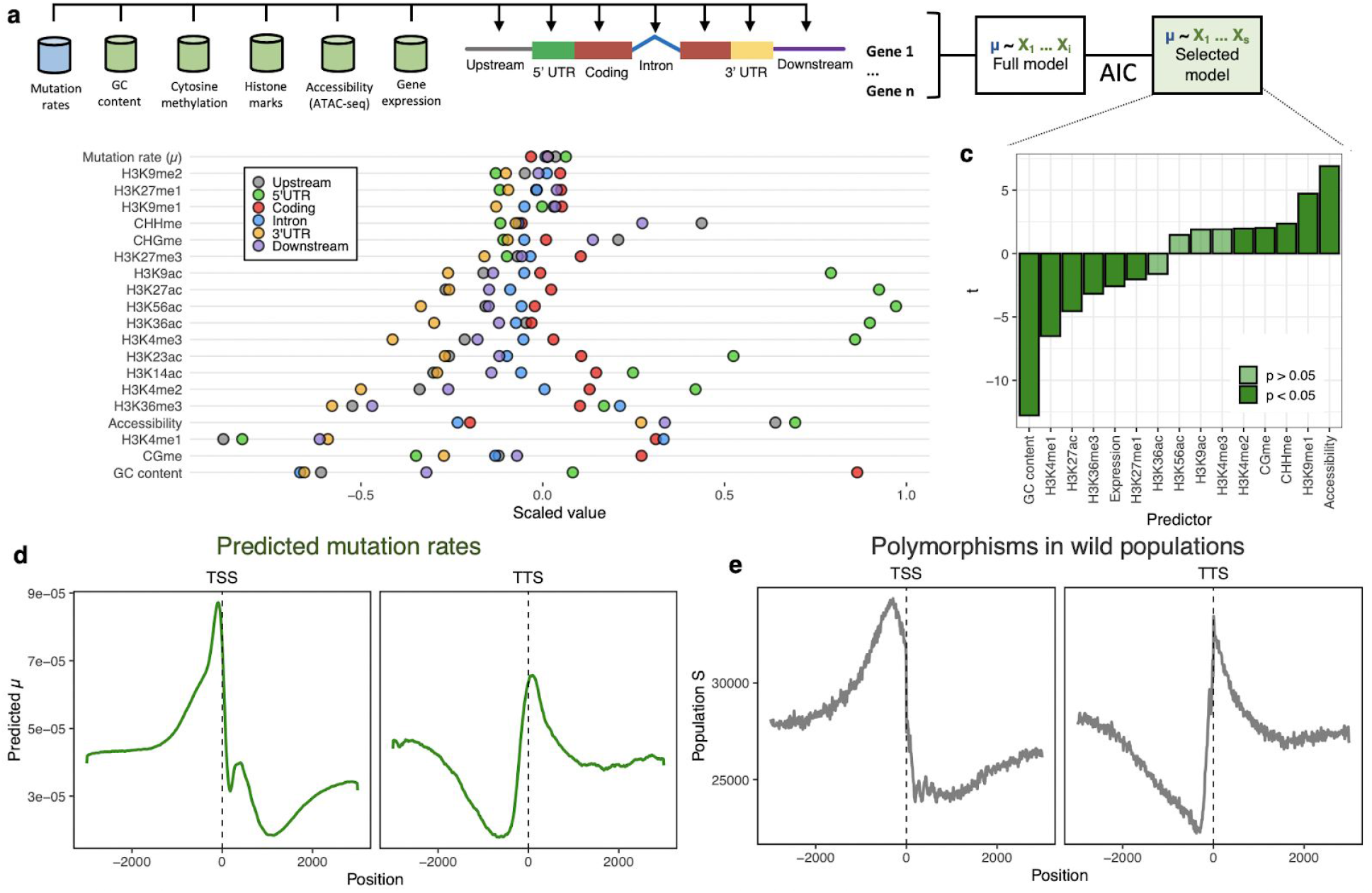
DNA sequence and epigenetic features predict mutation rates (# mutations per base pair) and explain distributions of natural polymorphisms around genes. (a) Graphical representation of modelling approach. (b) Variation in prevalence of epigenomic features across types of genic regions. Points show mean value for each epigenomic feature of each genic region across all genes. (c) Epigenomic features predict mutation rates. Optimum predictive model of mutation rates across genic regions based on AIC. Bars show the t-value of each predictor variable from the optimum generalized linear model. (d-e) Mutation rate variation drives levels of polymorphism such as reduced variation within gene bodies. (d) Predicted mutation rates based on optimum model and (e) observed number of segregating polymorphisms in natural populations (S) in 10 bp windows around transcription start (TSS) and transcription stop (TTS) sites. Note that these TSS and TTS plots are agnostic to gene length.

We then created a multiple linear regression model where the response variable was experimentally detected *de novo* mutation rate per base pair and predictor variables were scaled epigenomic features across each genic region and gene expression of each gene. To avoid overfitting caused by correlated predictor variables (Extended Data Fig. 5), we selected a model with the lowest Akaike information criterion (AIC) value (Fig. 2c).

The resulting model (Fig. 2c) included features previously linked to mutation rate. The observed negative correlation between GC content and mutation rate was consistent with other observations of lower mutation rate in GC rich regions ^23,39–42^ and mechanistically with both GC biased gene conversion ^43^ and lower rates of cytosine deamination in GC rich regions ^10,44–47^.

Furthermore, the histone modifications H3K4me1 and H3K27ac are known to be associated with lower mutation rates, particularly in active genes ^21,48–50^, and several studies have revealed explicit connections between H3K36me3 and DNA mismatch repair ^13,16,17^. These chromatin marks were enriched in gene bodies (Fig. 2b), and may help to explain the recent discovery that DNA mismatch repair preferentially targets genic regions - especially coding regions and introns - in *A. thaliana* ^22^. On the other hand, cytosine methylation, more prone to spontaneous cytosine deamination ^6,20,23,26,51^, was associated to high mutation rates, the heterochromatin-associated histone modification H3K9 methylation likely prevents repair machinery to be recruited and was associated to higher mutation rates ^6,20,23,26,51^, while highly accessible chromatin regions (such as those with active transcription factor binding sites) have previously been shown to interfere with nucleotide excision repair, which also led to higher mutation rates ^6,20,23,26,51^. Higher gene expression was associated with lower mutation rate, which differs from reported mutagenic effects of transcription in some organisms ^3,52^ but is consistent with transcription coupled repair and evidence of the tendency for DNA mismatch repair to preferentially target actively transcribed genes ^15,24,34,53^.

We tested the predictive performance of our model at a fine grained resolution by using it to predict per-bp mutation rates in 10 bp windows around transcription start and termination sites. Predicted mutation rates were considerably higher upstream and downstream of transcribed regions and lower within gene bodies (Fig. 2d, Extended Data Fig. 6). This pattern seemed to be primarily driven by reduced mutation rates in coding regions and introns (Extended Data Fig. 7) and could not be explained by variation in mapped read depth in these regions (Extended Data Fig. 8). We then calculated polymorphisms in natural accessions of *A. thaliana* in the same windows, which are known to be predominated by rare (i.e., evolutionarily young) variants ^54^ (Fig. 2e). The distribution of natural polymorphisms was positively correlated to the predicted mutation rates (r = 0.9, p<2×10^−16^), consistent with natural variation in genic regions being the product of mutation bias rather than only selection after mutation (Extended Data Fig. 7).

We also examined mutation rate variation within gene bodies. We observed variation in features that are predictive of mutation rate between gene exons (Extended Data. Fig. 9) and therefore calculated predicted mutation rates across coding region exons, grouping genes by their total number of coding exons. This revealed a gradient in predicted mutation rates with higher mutation rates in the extreme 5’ and 3’ coding exons (Fig. 3a). We found that levels of natural polymorphism show a similar pattern to predictions based on spontaneous *de novo* mutations (Fig. 3b).

**Figure 3.**
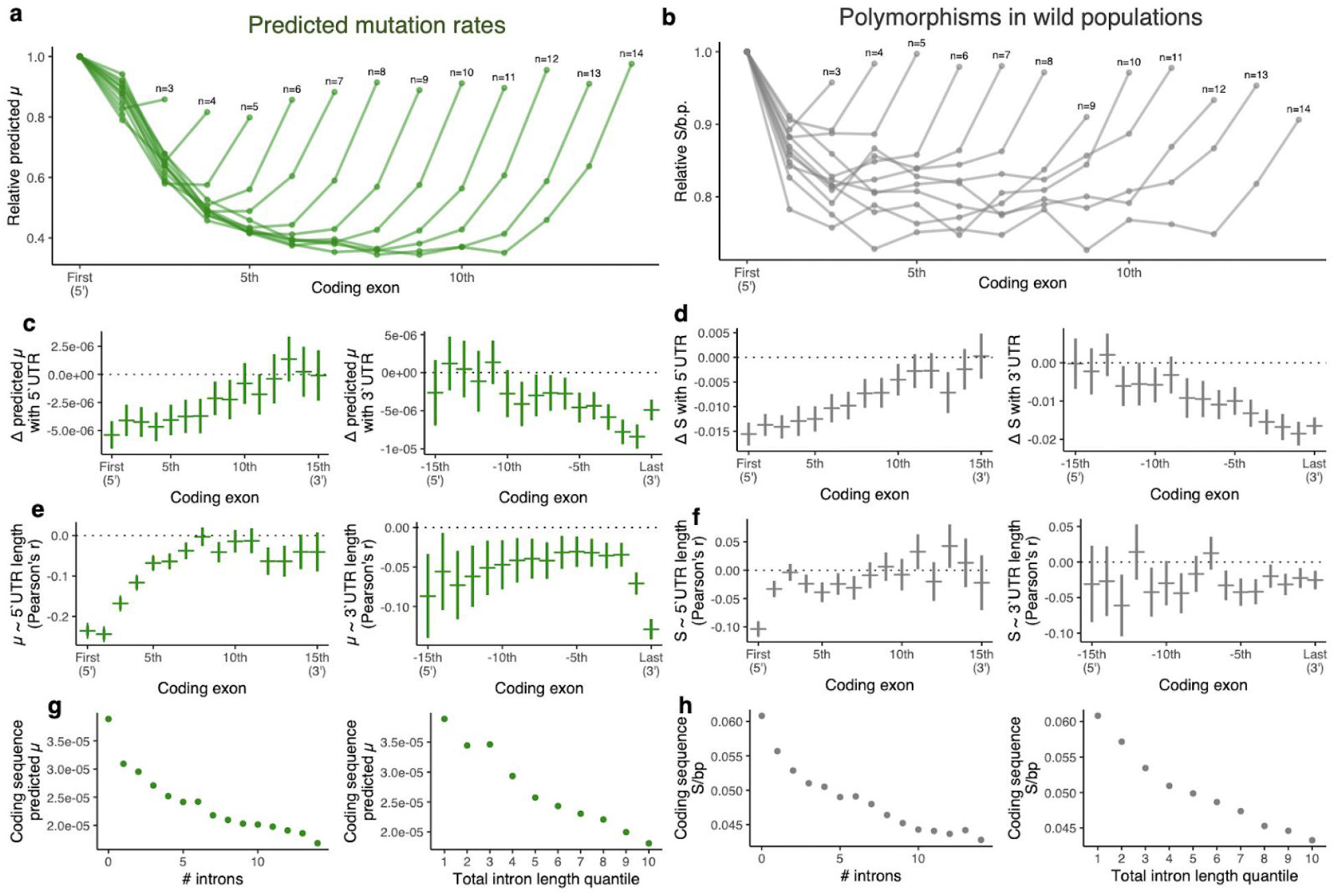
Introns and UTRs distance coding sequences from mutation hotspots. (a-b) Mutation and polymorphism rates are elevated at 5’ and 3’ extremes of transcribed sequences. (a) Relative predicted mutation rates (mutations per bp) and (b) levels of segregating polymorphisms (S) in sequential coding region exons for genes with different numbers of exons (n). Points and lines connect means for all genes with the same number of exons (n). (c-d) The presence of untranslated exons is associated with lower mutation rate in nearby coding regions. (c) Difference between mutation rates, and (d) levels of segregating polymorphism (S) between genes with and without untranslated regions (UTR). Left panels show the estimated effect of 5’ UTR and right panels show the effect of 3’ UTR. Horizontal lines mark the mean difference between gene sets (with and without UTR) for sequential exons in relation to the 5’ and 3’ end of the transcribed sequence. Vertical lines mark the confidence interval of a t-test. (e-f) For genes that do have UTRs, longer untranslated exons are associated with lower mutation rates in nearby coding regions. (e) Correlation between sequential exon mutation rates and (f) levels of polymorphism with the length in base pairs of 5’ (left panels) and 3’ (right panels) UTRs. Horizontal lines mark the Pearson correlation coefficient and vertical lines mark the confidence interval. (g) Number of introns and total intron length are negatively correlated with predicted mutation rates of coding regions. (h) Number of introns and total intron length are also negatively correlated with segregating polymorphisms in natural populations. Points indicate mean values.

The observation that mutation rates are elevated in leading and terminal coding exons suggests a cost to having coding regions near apparent mutation hotspots at the end of transcribed sequences in *A. thaliana*. This led us to hypothesize that untranslated regions (UTRs) and introns could provide a solution to this problem by distancing coding regions from the mutation prone sequences found at the extreme 5’ and 3’ ends of transcribed regions, which may also help explain the enrichment in introns of evolutionarily ancient and conserved genes ^55–57^. To test this, we compared predicted mutation rates in coding exons of genes with and without 5’ and 3’ UTRs. Predicted coding region mutation rates were 30.3% higher in genes annotated as lacking 5’ UTRs and 39.8% higher in genes lacking 3’UTRs. Consistent with a physical distancing effect, the inferred effect size of 5’ UTRs and 3’ UTRs on coding exon mutation rates was spatially non-random, being greatest in extreme 5’ and 3’ coding exons (Fig. 3c). Analyses of natural variation across these regions showed similar patterns (Fig. 3d). We also observed negative correlations between UTR lengths and coding exon mutation rates and natural polymorphisms, with the signal being strongest in the extreme 5’ and 3’ coding exons (Fig. 3e,f). To test the hypothesis that introns also buffer coding regions from mutation, we compared coding region mutation rates and the number and length of introns. Predicted coding region mutation rates were 90.7% higher in genes lacking introns. Predicted mutation rates in coding regions were lower in genes with greater intron number (r = −0.37) and length (r = −0.34) (Fig. 3g). These patterns were mirrored by levels of natural polymorphisms (Fig. 3h).

The preceding analyses indicated that epigenomic features such as histone modifications and gene expression and sequence features such as the existence of introns and UTRs provide explanatory mechanisms of mutation rate variation between genes (Fig. 4a,b). To test whether mutation rate variation is driving rates of gene coding sequence evolution, we first evaluated the relationship between predicted mutation rates and putatively neutral variation in wild *A. thaliana*. These analyses confirmed that predicted mutation rates were positively correlated with rates of synonymous molecular polymorphism in *A. thaliana* populations and divergence from *A. lyrata* (Fig. 4c, d), thus supporting the notion that genome-wide mutational bias largely shapes gene evolution (Fig. 4c, d).

**Figure 4.**
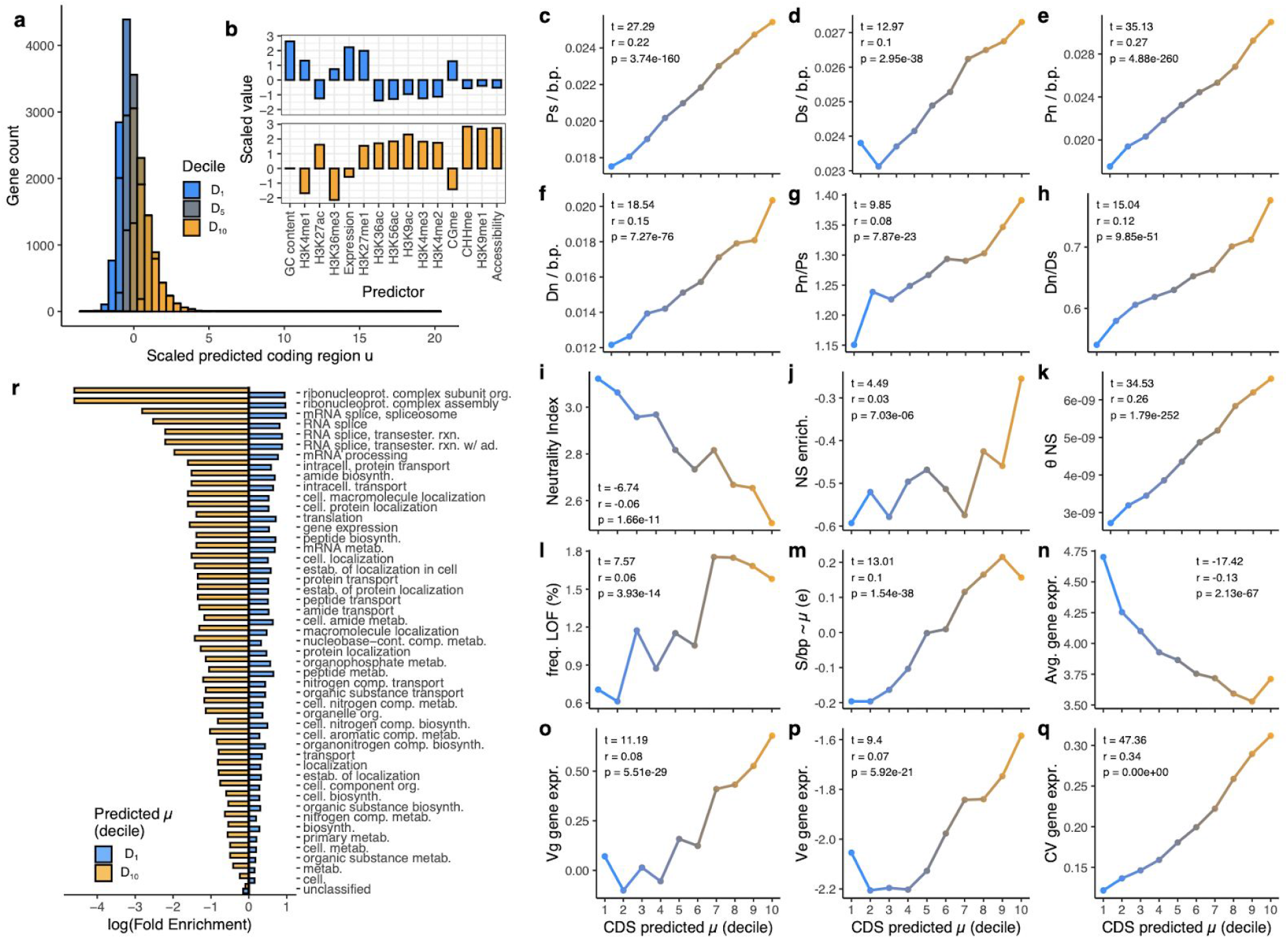
Functionally constrained genes have lower predicted mutation rates. (a) Distribution of predicted mutation rates in coding regions across genes. (b) Cytogenetic features predictive of mutation rate between top and bottom deciles of predicted mutation rate. Relationships between predicted coding sequence (CDS) mutation rates and (c) levels of synonymous polymorphism (Ps) per base pair in natural populations, (d) levels of synonymous divergence (Dn) per base pair from *A. lyrata*, (e) levels of non-synonymous polymorphism (Pn) per bp in natural populations, (f) levels of non-synonymous divergence (Dn) per bp from *A. lyrata*, (g) ratio of Pn to Ps, (h) ratio of Dn to Ds, (i) neutrality index ((Pn/Ps)/(Dn/Ds)), (j) enrichment of non-synonymous polymorphisms compared to genome-wide average, (k) Watterson’s diversity measure for non-synonymous variants, (l) frequency of loss-of-function (LOF) alleles observed in natural populations, (m) scaled residuals of linear predictive model of polymorphism in relation to mutation rate, (n) mean gene expression level across natural populations, (o) genetic variance of gene expression across natural populations, (p) environmental variance of gene expression across natural populations, and (q) coefficient of variation (CV) in gene expression across natural populations. Points on each plot mark the mean value of genes grouped into deciles according to predicted mutation rates. Colors points and connecting lines reflect these quantiles, as shown in (a). Summary statistics of correlation between gene level predicted mutation rates and other variables are displayed in each plot. (r) Gene ontology (GO) terms enriched in genes from bottom and top deciles of coding region mutation rates.

So far, these results were compatible with neutral variation in mutation rates. To finally test the hypothesis that gene level mutation rates are non-random with respect to likely fitness consequences of mutations between genes, we compared predicted coding region mutation rates between genes in *A. thaliana* with various estimates of evolutionary and functional constraint. We observed significant correlations between predicted mutation rates and all estimates of functional constraint (Fig. 4e-q), including estimates of constraint on sequence (Fig. 4e-m) and regulatory function (Fig. 4n-q). We attempted to disentangle the importance of selection and mutation rate variation shaping levels of natural polymorphisms by calculating the residuals between the expected level of polymorphisms from the DNA sequence and epigenetic features, and the observed levels of natural polymorphisms (Fig. 4m). This analysis showed that in genes where mutation rate is predicted to be low, the observed number of polymorphisms tends to be even lower than that predicted by mutation rate alone, which is consistent with the action of purifying selection on natural polymorphisms rather than on the predicted mutation rates. In addition to being consistent with the other analyses that showed lower mutation rates in genes that experience greater purifying selection, this result also provided further evidence that selection on the mutation accumulation lines used here is unlikely to explain these results. When looking at the top and bottom decile of genes in terms of predicted coding region mutation rates (Fig. 4r), we found that they differ according to biological function. While annotations of genes with the lowest predicted mutation rates were enriched for core (ie. housekeeping) biological functions, genes with the highest predicted mutation rates tended to be significantly depleted for these same functions (Fig. 4r).

While further functional and modeling studies potentially incorporating complex interactions between demography, selection, and mutation variation will be required to more fully separate these effects, it seems clear that variation in cytogenetic states explaining mutation rate biases is not independent from gene function, contradicting the classical expectation that mutation rates are random with respect to the fitness consequences. Instead these results support mutation rates being lower in functionally constrained genes where mutations are more likely to be deleterious ^5,7,8^.

Evolutionary theory predicts that beneficial gene level mutation rates could readily evolve if ΔU * L_segment_ * N_e_ > 1 (where ΔU = reduction in deleterious mutation rate, L_segment_ = length of sequence affected, N_e_ = effective population size) ^3,8^. This criterion is met if processes governing mutation rates interact with cytogenetic regulatory features to preferentially target multiple important (effectively large ΔU) genes – resulting in a large effective L_segment_ – for repair ^3,8^. Consistent with previous functional work, here we find such features are indeed predictive of mutation rates and are distributed non-randomly between genes according to function (functionally constrained genes are enriched for features that are linked to lower mutation rates, Fig. 4, Extended Data Fig 10). In contrast to other models of beneficial mutation rate evolution that invoke gene-specific modifiers of mutation rate, this scenario of genic mutation rate evolution requires that selection is sufficient to maintain gene level regulatory features, not gene level mutation rates directly. The observations made in this investigation are thus consistent with a growing body of research suggesting that this model of beneficial genic mutation rate evolution is both theoretically and empirically plausible.

The implications of these findings for evolutionary biology are far-reaching. If mutation rates are specifically lower in functionally constrained genes, one might hypothesize that the distribution of fitness effects of new mutations would be skewed and adaptive evolution would proceed faster than predicted from models assuming that mutation probabilities are truly independent of mutational consequences. We thus believe that the correlated effect of natural selection and variable mutation rate provides a more complete explanation of natural genetic variation and gene evolution in *A. thaliana*.

## Extended Data

**Extended Data Figure 1.**
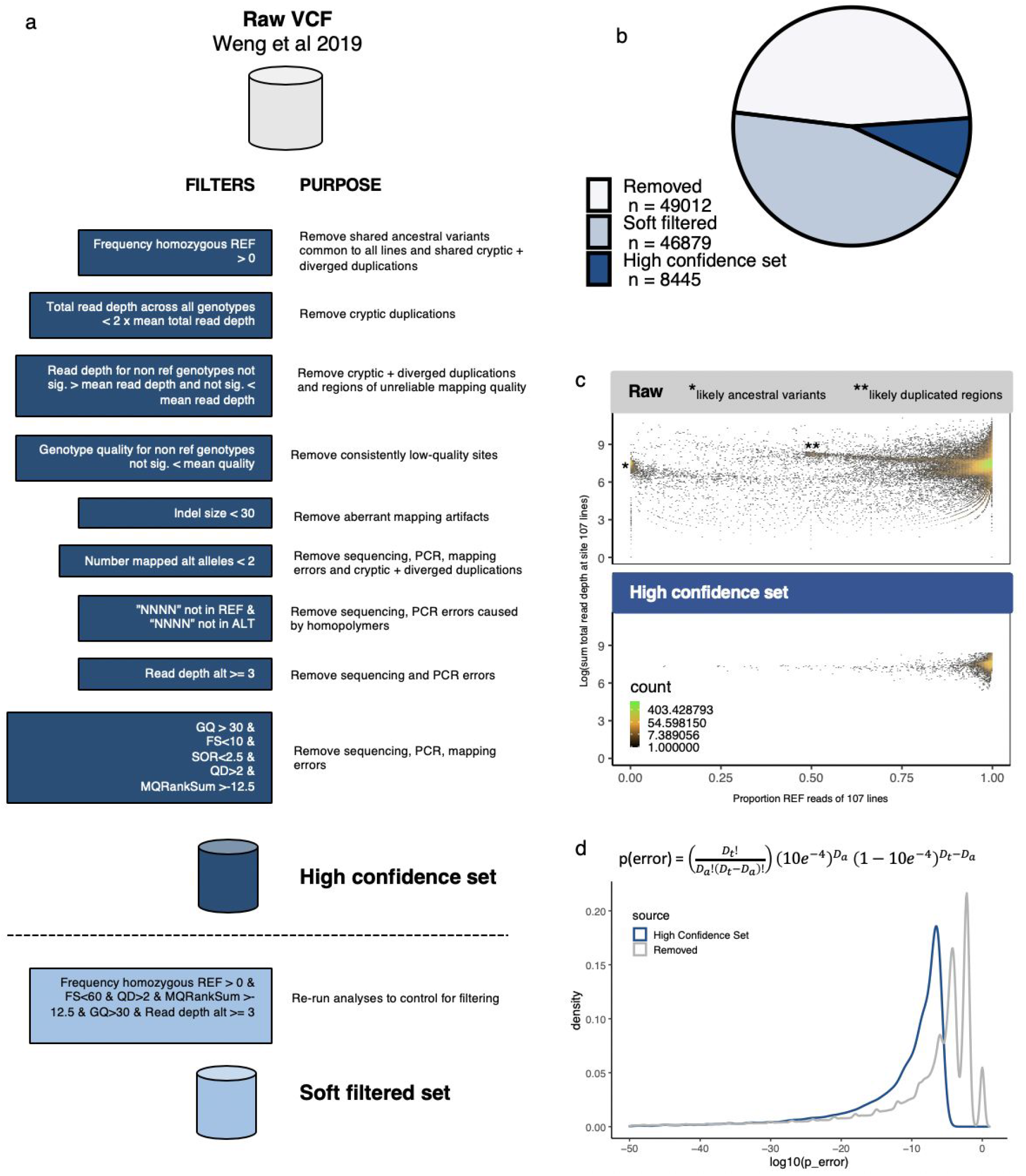
Workflow and quality control of *de novo* mutation identification. (a) Filtering pipeline. (b) High-quality *de novo* mutations called in this investigation (total number = mutations here + Weng et al. 2019.) (c) Visualization of raw data and high confidence set. (d) Estimate probability of call being error based on alternative and total read depths in the high confidence set and variants removed by filtering.

**Extended Data Figure 2.**
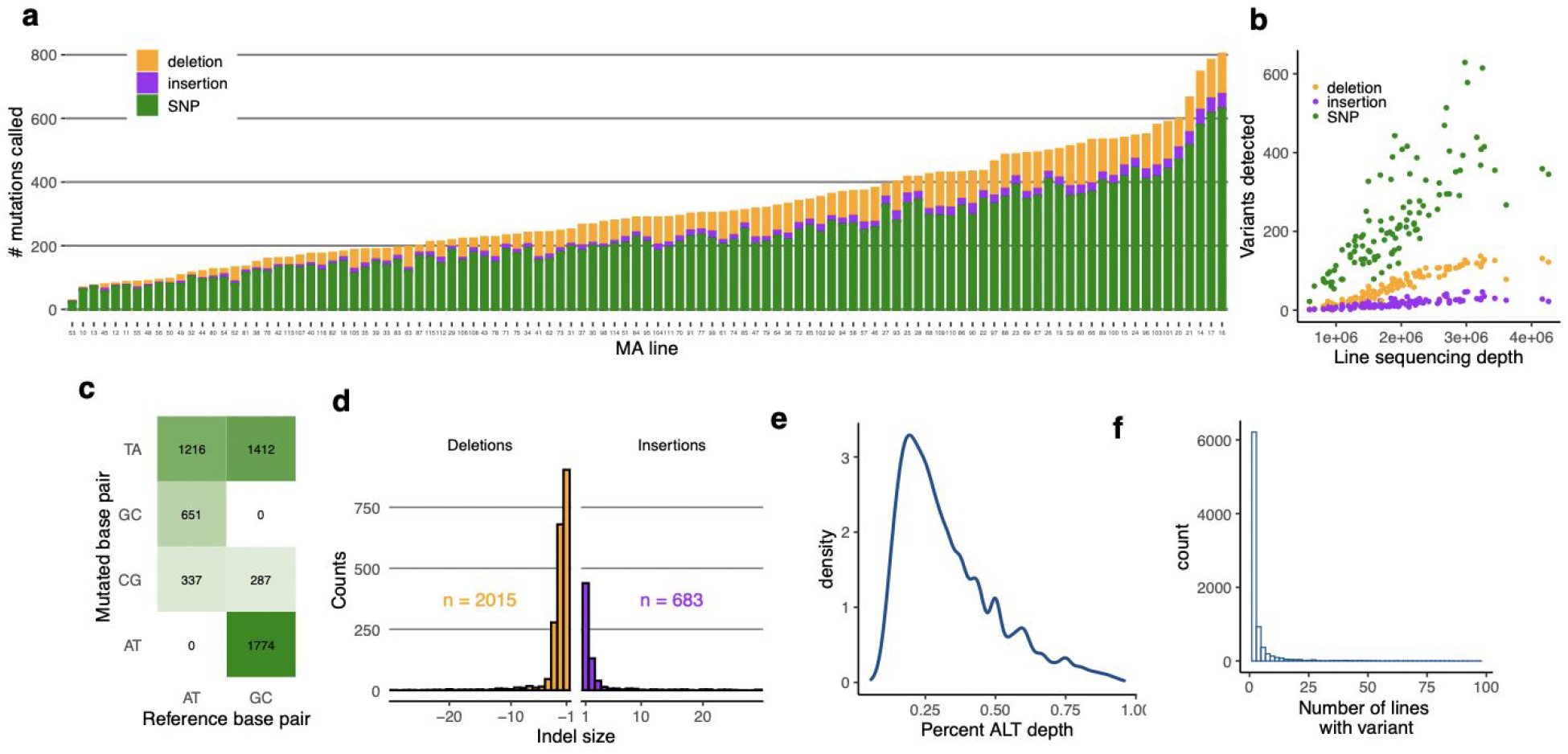
Summary of *de novo* mutations called in this study. (a) Number of mutations detected in each MA line. (b) Relationship between number of mutations detected and total sequencing depth in MA lines. (c) Frequencies of single nucleotide transitions and transversions. (d) Size distribution of insertions and deletions. (e) Distribution of alternative allele read depth for putative somatic mutations (f) Distribution of frequency of specific mutations across lines.

**Extended Data Figure 3.**
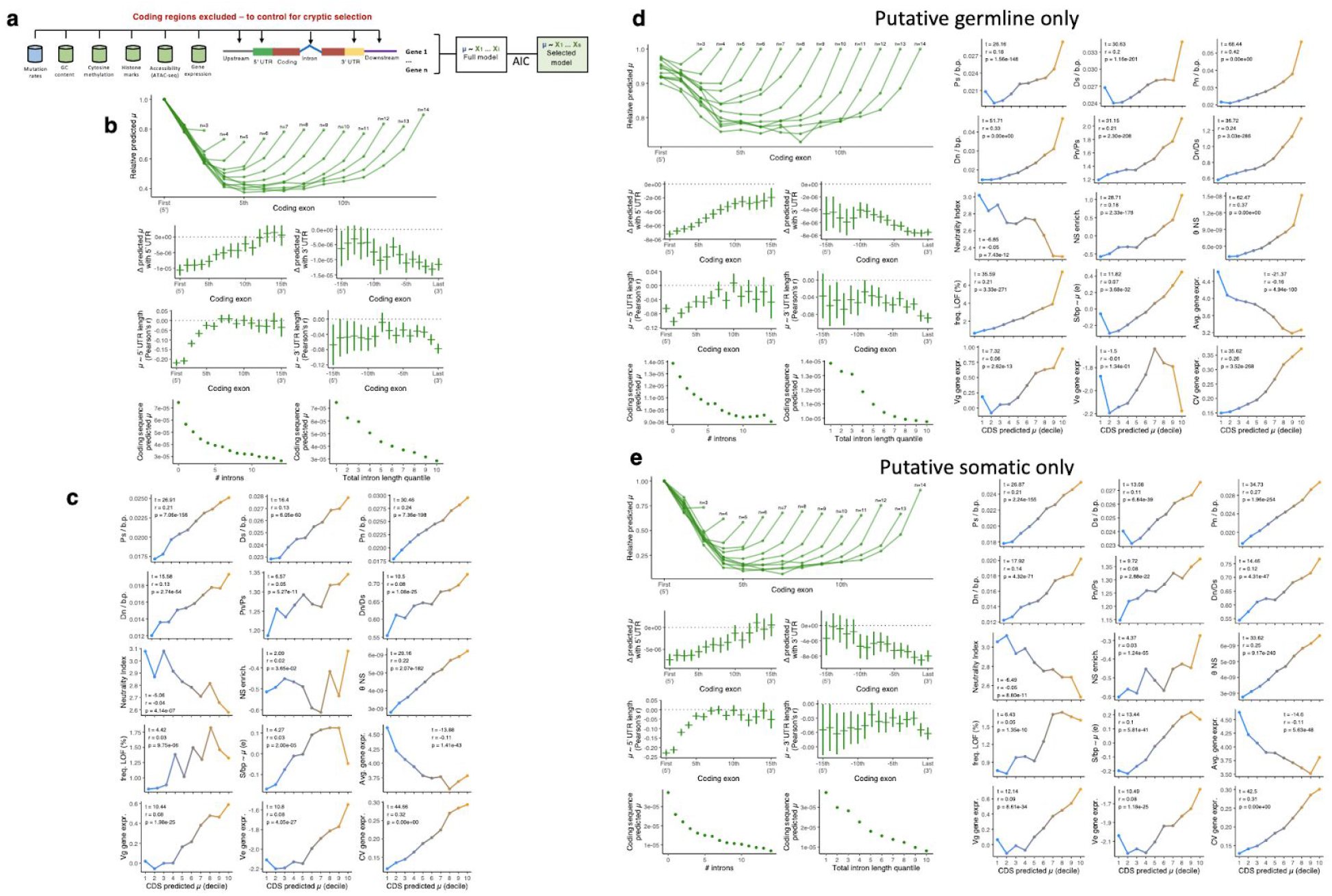
Schematic and example results from analyses based on models where coding regions were excluded in training the model and analyses of only putative germline or putative somatic mutations. Conclusions drawn from these results are qualitatively the same as those drawn from results in the main text. This provided support that main results are not an artifact of selection on coding regions interfering with ability to create an unbiased estimate of genome-wide mutation processes. (a) Schematic of modeling without coding regions. (b) Corresponds to Fig. 3, and (c) Fig. 4 results. (d) Aanalyses run with putative germline mutations only. (e) Aanalyses run with putative somatic mutations only.

**Extended Data Figure 4.**
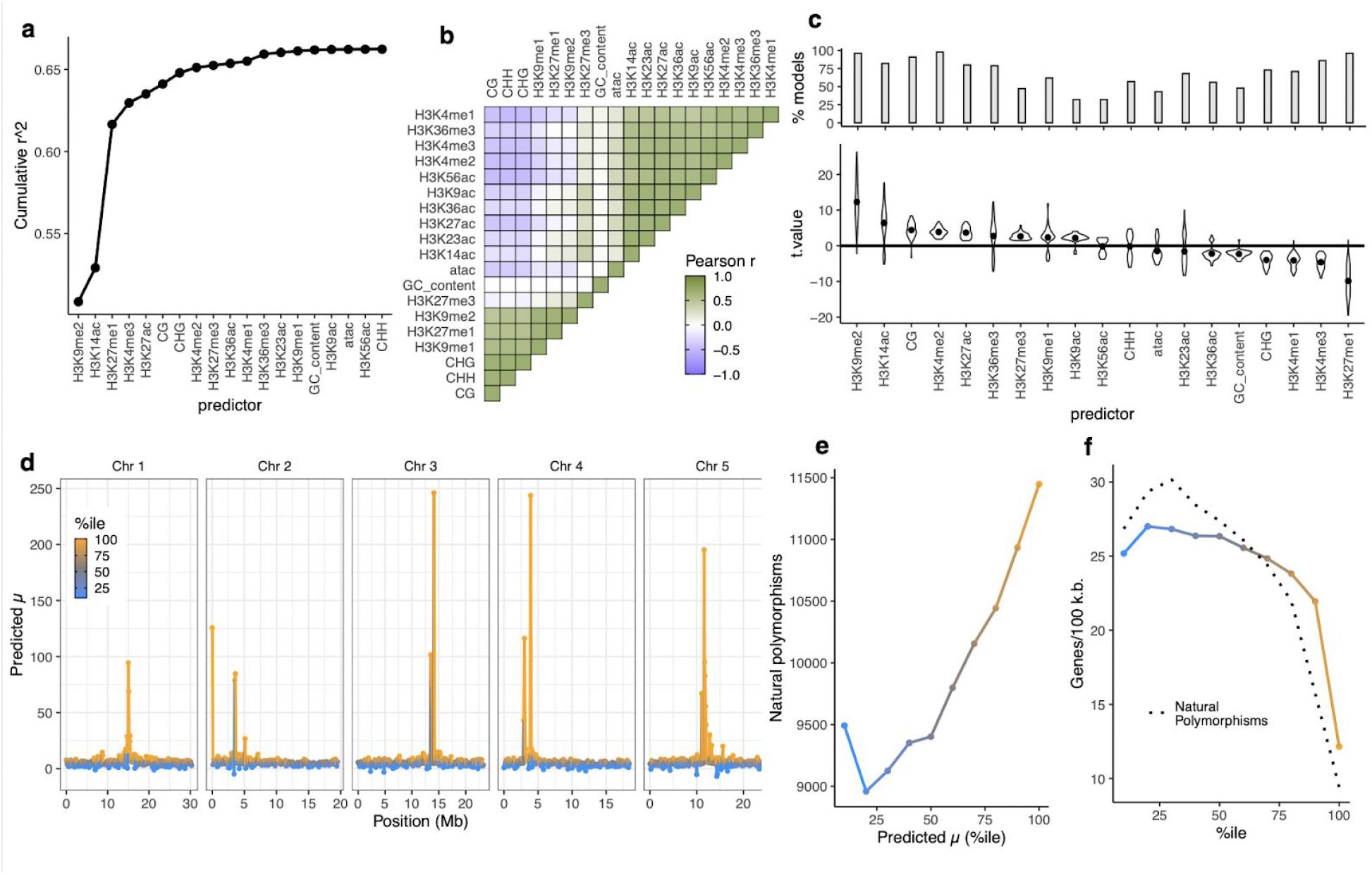
Epigenomic features predict mutation rate variation at regional resolution (100 kb windows). (a) Cumulative explanatory power of predictor variables added sequentially by forward model selection. (b) Correlations between predictor variables (c) Summary from 100 iterations of 50:50 cross validation where the best model was selected by AIC. Top panel shows fraction of selected models which included that predictor. Bottom panel shows the distribution of t-values of each predictor when included in the selected model. (d) Predicted mutation rates in test sets from 100 iterations of 50:50 cross validation. (e) Relationship between mean predicted mutation rates in test sets with levels of natural polymorphism. (f) Relationships between predicted mutation rates and natural polymorphism with gene content.

**Extended Data Figure. 5.**
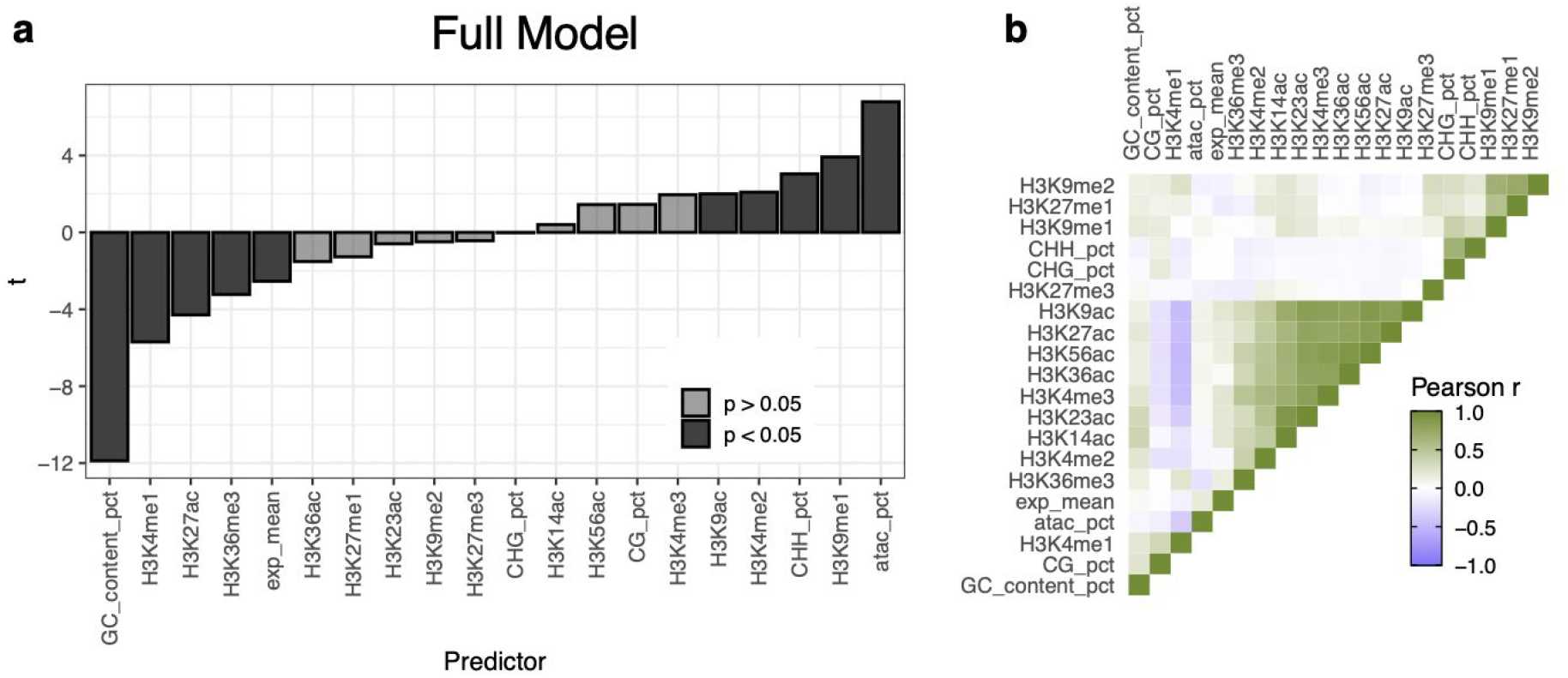
(a) Summary statistics of full multiple linear regression model model of u ~ epigenomic features for genic regions (upstream, UTR, coding, intron, downstream) before selecting a limited model by AIC. (b) Correlations between predictors.

**Extended Data Fig.6.**
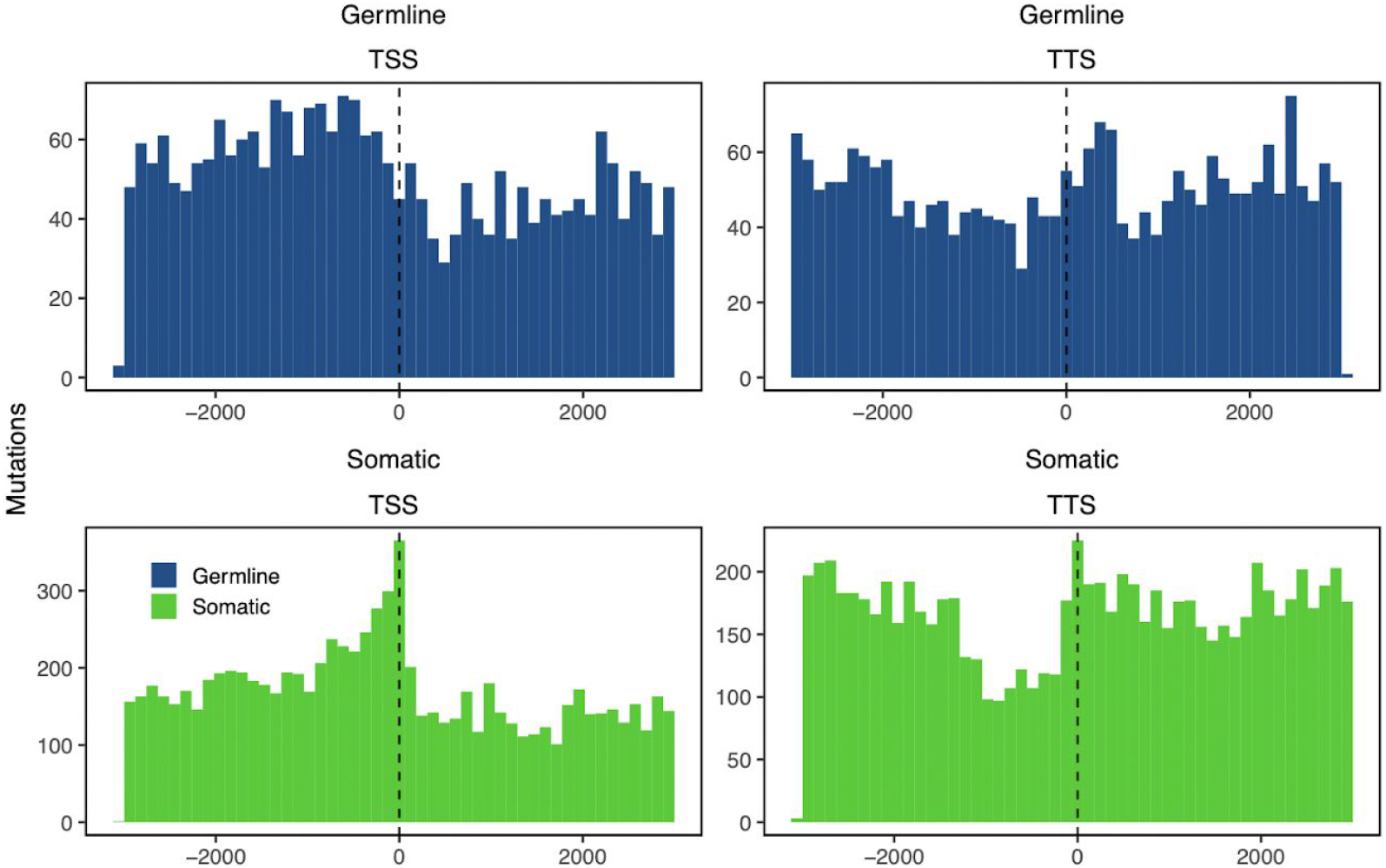
Raw numbers of *de novo* mutations detected around TTSs and TSSs.

**Extended Data Figure 7.**
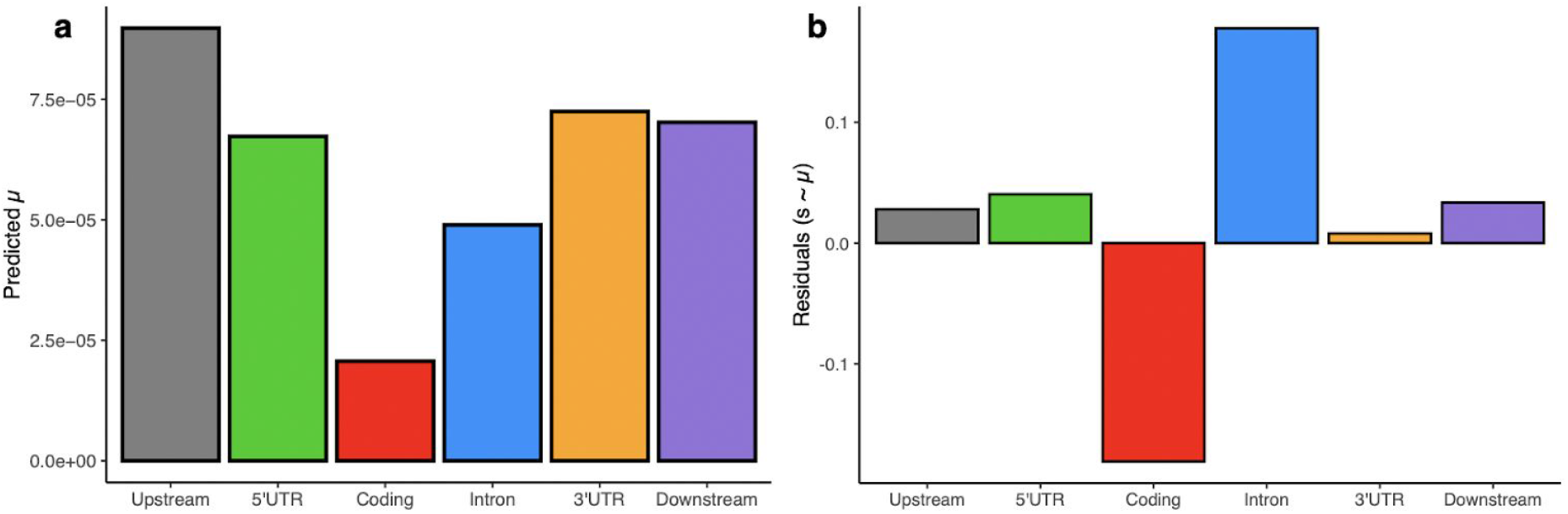
(a) Predicted mutation rates, and (b) scaled residuals ((Obs-Pred)/Pred) from S ~ u. Significantly negative residuals in coding regions are consistent with purifying selection in natural populations acting on new mutations.

**Extended Data Figure 8.**
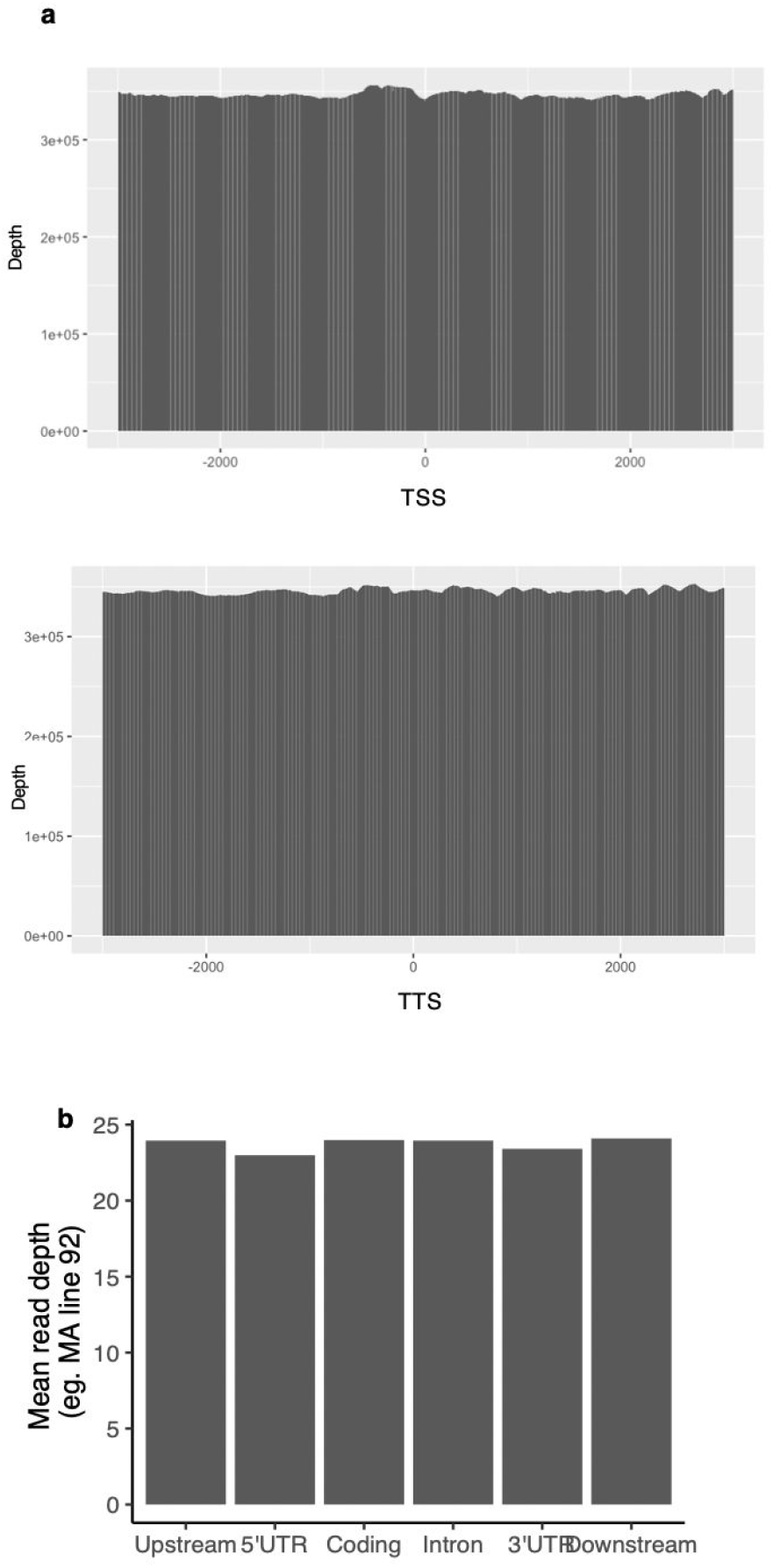
Sequencing depth does not correlate with natural polymorphisms or mutation distributions. Example total sequencing depth around (a) TSS and TTS, and (b) mean sequencing depth in different genic regions.

**Extended Data Figure 9.**
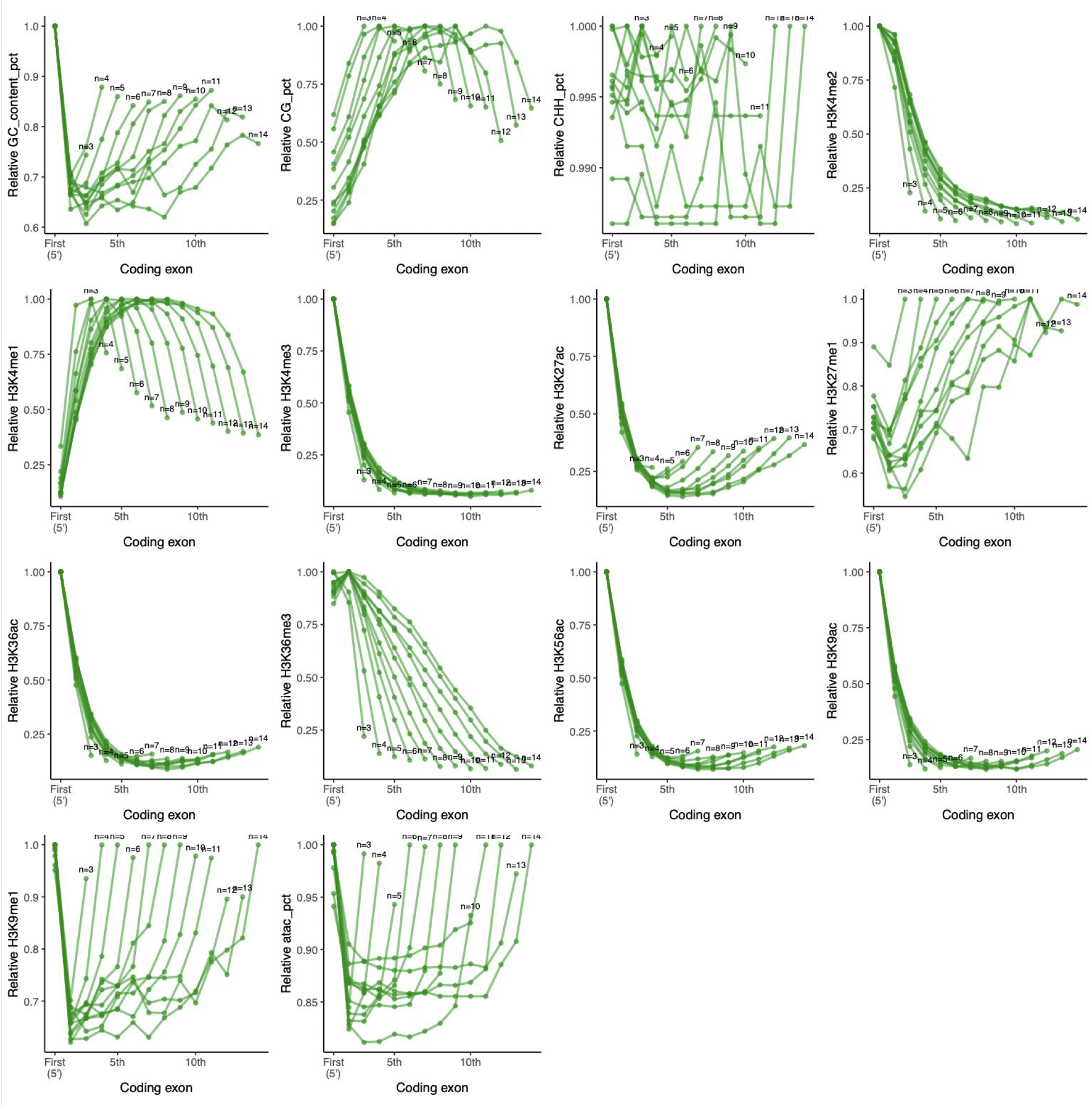
DNA sequence and epigenomic features (from selected AIC model, Fig. 2 C) across genes of varying exon number (shown are 2 < n < 15).

**Extended Data Figure 10.**
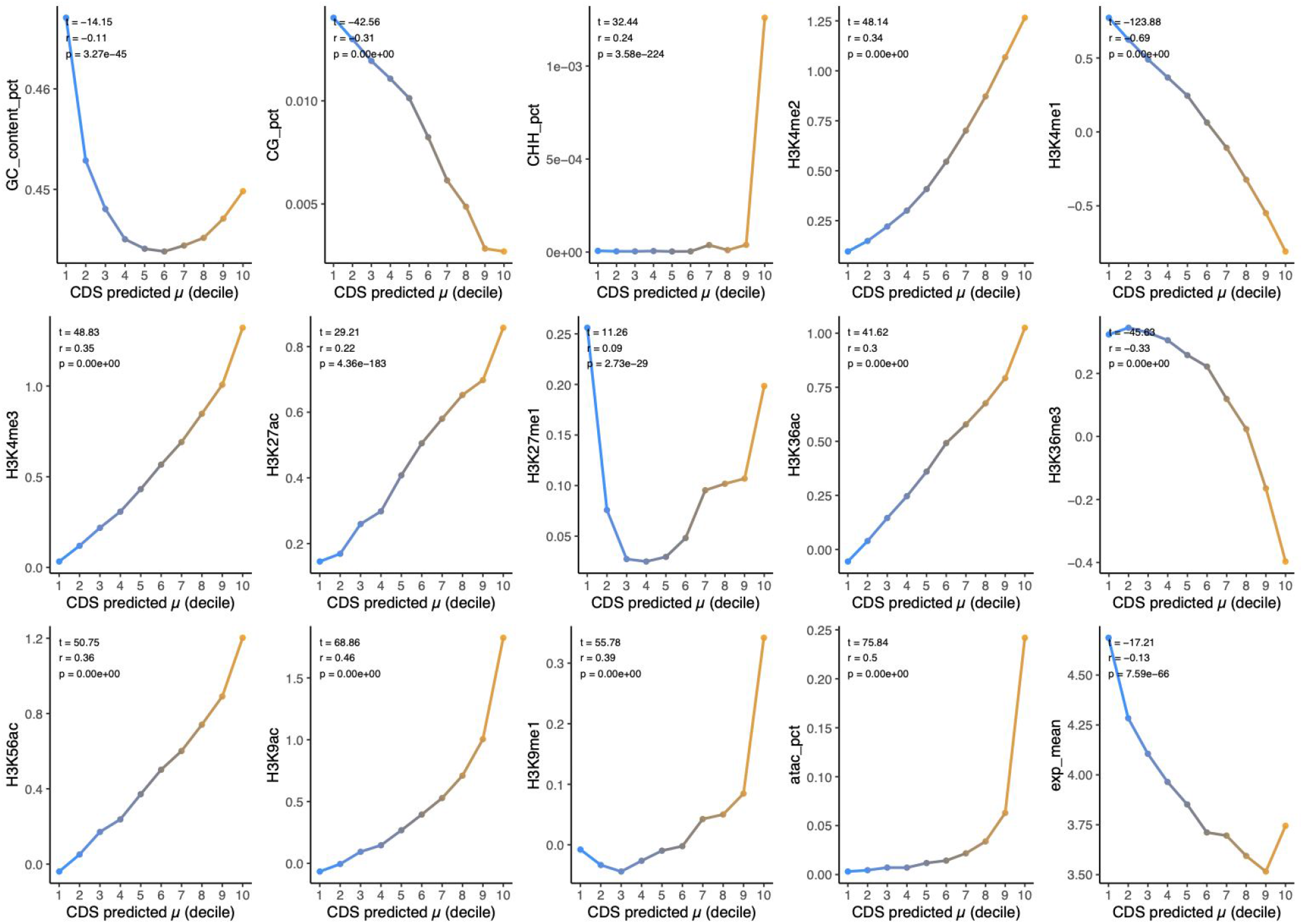
Epigenomic features in genes ranked into deciles by predicted mutation rates.

## Acknowledgements

We thank Bob Schmitz, Dan Sloan, Jeff Ross-Ibarra, Mike Lynch, Dmitri Petrov, Padraic Flood, Dan Runcie, Ksenia Krasileva, and members of the Weigel Lab for insightful conversations and valuable comments on earlier versions of this manuscript. This work was supported by the Max Planck Society.

## Materials and Methods

### Identification of *de novo* mutations in *A. thaliana*

Mutations were identified from 107 mutation accumulation lines of the *A. thaliana* Col-0 accession, which is the basis of the *A. thaliana* reference genome ^6^. The lines had been previously grown for 24 generations of single-seed descent before pooled Illumina sequencing (100 bp paired-end reads) of approximately 40 seedlings at the 4-leaf stage (2-weeks old) of the 25^th^ generation. Variants were called using GATK HaploCaller ^6^. Whereas in humans, germline mutations are primarily influenced by processes specific to reproductive organs ^25^, because plants lack a completely segregated germline ^38^, we hypothesized that mechanisms which influence local mutation rates in the germline may be reflected in the distribution of somatic mutations as well. In addition to the original variants called, we implemented a custom filtering pipeline to identify a high confidence set of additional *de novo* mutations (Extended Data Fig. 1). This set included somatic variants and additional germline variants that were not called in the original analyses of these mutation accumulation lines. Somatic mutations were previously excluded because they appear as heterozygous calls. Germline mutations were previously excluded if at least 1 out of the 107 lines also included a putative somatic mutation at the same position. Based on previously described per generation germline mutation rate (1-2) and with the knowledge that these lines were self fertilized each generation, we expect the seedlings which were sequenced to be segregating for 2-4 heterozygous germline variants, which would have been called as somatic mutations by our pipeline (approximately 2-5% of putatively somatic mutations). Because we combined putative somatic and germline mutations to characterize the mutational landscape of the *A. thaliana* genome, this did not have an obvious effect on our results.

### Estimating selection on de novo mutations

To estimate the effect of selection on *de novo* mutations found here, we examined rates of synonymous, non-synonymous, and stop-gained variants-which provide a robust estimate of selection because they are all of the same mutational class (SNPs) and are not biased by true mutation rate variation (i.e., coding region vs. introns). As a basis of comparison we also calculated the rates of synonymous, non-synonymous and stop-gained SNPs in natural populations of *A. thaliana*, which are subject to long-term natural selection. We also derived an expected null ratio of nonsynonymous to synonymous mutations using knowledge on the relative base composition of all coding regions in the reference genome, the relative proportion of coding region mutations (e.g., C->T are most common), and the proportion of all possible codon transitions that lead to synonymous vs nonsynonymous mutations. Ratios of non-synonymous to synonymous and stop gained to synonymous were compared between *de novo* mutations and those observed in natural populations and the null expectation by chi-squared tests.

### Sequence and epigenomic features

To build a high resolution predictive model of mutation rate variation across the *A. thaliana* genome we extracted or generated data describing sequence and epigenomic features across the genome. First, we calculated GC content (% of sequence) across regions. We also downloaded from the Plant Chromatin State Database 62 BigWig formatted datasets characterizing the distribution of histone marks ^58^. For each specific histone mark, depths were scaled and averaged across each region for downstream analyses.

### Col-0 cytosine methylation

Methylated cytosine positions in the *A.thaliana* Col-0 (6909) wild-type leaf methylome were obtained from the 1001 Genomes and Epigenomes dataset GSM1085222 ^59^ under the file GSM1085222_mC_calls_Col_0.tsv.gz. Cytosines were further classified into three categories (CG/CHG/CHH) for all downstream analyses. For each region we calculated the number of methylated cytosines in each category per bp.

### Plant growth and conditions for ATAC-seq

Seedlings of Col-0 genotype were cold treated (−80°C), ethanol-sterilized, and stratified in 0.1% Agar, on MS-Agar (+Sucrose) plates at 4°C for 4 days in the dark. Plates were then kept vertical in 23°C long-day Percival Chambers. On the 11^th^ day of light exposure, 10-20 seedlings each from three MS-Agar plates were fixed with formaldehyde by vacuum infiltration and stored at −80°C before nuclei extraction and ATAC-seq.

### Nuclei extraction

Fixed tissue was chopped finely with 500 μl of General Purpose buffer (GPB; 0.5 mM spermine•4HCl, 30 mM sodium citrate, 20 mM MOPS, 80 mM KCl, 20 mM NaCl, pH 7.0, and sterile filtered with 0.2 μm filter, followed by the addition of 0.5% of Triton-X-100 before usage). The slurry was filtered through one-layered Miracloth (pore size: 22-25 μm), followed by filtration through a cell-strainer (pore size: 40 μm) to collect nuclei. Approximately 50,000 DAPI stained nuclei were sorted using fluorescence-activated cell sorting (FACS) as two technical replicates.

### ATAC-seq library prep

Sorted nuclei were heated at 60°C for 5 minutes, followed by centrifugation at 4°C (1,000 g, 5 minutes). Supernatant was removed, and the nuclei were resuspended with a transposition mix (homemade Tn5 transposase, a TAPS-DMF buffer and water) followed by a 37°C treatment for 30 minutes. 200 μl SDS buffer and 8 μl 5 M NaCl were added to the reaction mixture, followed by 65°C treatment overnight. Nuclear fragments were then cleaned up using Zymo PCR column-purification (DNA Clean and Concentrator) columns. 2 μl of eluted DNA was then subjected to 13 PCR cycles, incorporating Illumina indices, followed by a 1.8:1 ratio clean-up using SPRI beads. Genomic DNA libraries were prepared using the same library prep protocol from the Tn5 enzymatic digestion step onwards.

### Sequencing, read alignment and peak calling

Libraries were sequenced in paired-end mode, using an Illumina HiSeq 3000 instrument. Each technical replicate (derived from nuclei sorting) was sequenced at a depth of 3.5 million paired end reads. The reads were aligned as two single-end files to the TAIR10 reference genome using *bowtie2* [default options], filtered for the SAM flags 0 and 16 (only reads mapped uniquely to the forward and reverse strands), converted separately to bam files. The bam files were then merged, sorted, and PCR duplicates were removed using *picardtools.* The sorted bam files were then merged with the corresponding sorted bam file of a second technical replicate (samtools merge --default options) to get a final depth of approximately 6 million reads for each biological replicate.

Peak calling was carried out for each biological replicate using *MACS2* using the following parameters:

~~~
macs2 callpeak -t [ATACseqlibrary].bam -c [Control_library].bam -f BAM --nomodel --shift -50 --extsize 100 --keep-dup=1 -g 1.35e8 -n [Output_Peaks] -B -q 0.05
~~~

Peak files and .bam alignment files from three biological replicates were then processed with the R package DiffBind to identify consensus peaks which overlapped in at least two out of three peaksets (FDR <0.01). The library quality was estimated by measuring the FRIP (Frequency of reads in peaks) scores for the three replicates, which were 0.36,0.36 and 0.39 (above the standard quality threshold of 0.3). These consensus peaks were used for all further downstream analyses, such as intersection with genic features and cytosine methylation positions.

### Gene expression

Gene expression was calculated at each locus by the mean expression detected across 1,135 accessions ^59^. Furthermore, we used these same data to extract the genetic variance (Vg) and environmental variance (Ve) in expression levels for each gene. Finally, we calculated the coefficient of variation (variance/mean) for each gene.

### Predictive model of mutation rates

To create a genome-wide gene-level predictive model of mutation rate we created a generalized linear model where the response variable was detected mutation rate across every genic feature (upstream, UTR, coding, intron, downstream) and the predictor variables were GC content, classes of cytosine methylation, histone modifications, and expression of each gene. From this full model, a limited predictive model was selected based on forward and backward selection with the lowest AIC value by the stepAIC function in R. For downstream analyses we used the predicted mutation rates based on this model for genes, exons, and regions of interest from the TAIR10 genome annotation.

### Signatures of selection and constraint from natural populations

Gene level summary statistics for signatures of selection and constraint. Synonymous and non-synonymous polymorphism among natural *A. thaliana* accessions and divergence from A. lyrata (Pn, Ps, Dn, Ds, respectively) were calculated using mkTest.rb

(https://github.com/kern-lab/). The Neutrality Index developed by McDonald and Kreitman ^60^ was calculated from these values for each gene where data was available (not all genes have called orthologs in *A. lyrata*) as (Pn/Ps)/(Dn/Ds). Higher values of the Neutrality Index are traditionally interpreted as evidence of stronger purifying selection because non-synonymous variants in genes with such values tend not to become fixed. Enrichment of non-synonymous variants compared to genome wide average and Waterson’s diversity estimate (theta) of non-synonymous variation were calculated independently. The frequency of loss-of-function was calculated similar to previous descriptions ^61,62^, where loss-of-function was defined as alleles (premature stop codons and frameshifts) resulting in disruption of at least 10% of the coding region of the canonical gene model. Genes experiencing purifying selection should exhibit lower levels of natural polymorphism than what would be predicted by mutation rate alone. To test this, we built a linear model of coding region polymorphisms as a function of predicted mutation rates. We then calculated scaled residuals for each gene, and tested whether they are more negative in genes expected to be under purifying selection. To estimate constraint on gene regulatory function we looked at average expression across diverse genotypes. We also tested for relationships between predicted mutation rates and the coefficient of variation in gene expression, additive genetic variance for gene expression across diverse genotypes, and environmental variance in gene expression ^59^. To compare the biological function of genes according to predicted mutation rates we also analyzed gene ontology categories for genes in the top and bottom deciles ranked by predicted mutation rates^63^.

## References

1. Luria, S. E. & Delbrück, M. Mutations of Bacteria from Virus Sensitivity to Virus Resistance. Genetics 28, 491–511 (1943).

2. Loewe, L. Genetic mutation. Nature education 1, 113 (2008).

3. Lynch, M. et al. Genetic drift, selection and the evolution of the mutation rate. Nat. Rev. Genet. 17, 704–714 (2016).

4. Svensson, E. I. & Berger, D. The Role of Mutation Bias in Adaptive Evolution. Trends Ecol. Evol. 34, 422–434 (2019).

5. Martincorena, I., Seshasayee, A. S. N. & Luscombe, N. M. Evidence of non-random mutation rates suggests an evolutionary risk management strategy. Nature 485, 95–98 (2012).

6. Weng, M.-L. et al. Fine-Grained Analysis of Spontaneous Mutation Spectrum and Frequency in Arabidopsis thaliana. Genetics 211, 703–714 (2019).

7. Martincorena, I. & Luscombe, N. M. Response to No gene-specific optimization of mutation rate in Escherichia coli. arXiv [q-bio.GN] (2013).

8. Martincorena, I. & Luscombe, N. M. Non-random mutation: the evolution of targeted hypermutation and hypomutation. Bioessays 35, 123–130 (2013).

9. Stoletzki, N. & Eyre-Walker, A. The Positive Correlation between d N/d S and d S in Mammals Is Due to Runs of Adjacent Substitutions. Mol. Biol. Evol. 28, 1371–1380 (2011).

10. Hodgkinson, A. & Eyre-Walker, A. Variation in the mutation rate across mammalian genomes. Nat. Rev. Genet. 12, 756–766 (2011).

11. Terekhanova, N. V., Seplyarskiy, V. B., Soldatov, R. A. & Bazykin, G. A. Evolution of Local Mutation Rate and Its Determinants. Mol. Biol. Evol. 34, 1100–1109 (2017).

12. Chen, X. & Zhang, J. No gene-specific optimization of mutation rate in Escherichia coli. Mol. Biol. Evol. 30, 1559–1562 (2013).

13. Fang, J. et al. Cancer-driving H3G34V/R/D mutations block H3K36 methylation and H3K36me3-MutSα interaction. Proc. Natl. Acad. Sci. U. S. A. 115, 9598–9603 (2018).

14. Li, C. & Luscombe, N. M. Nucleosome positioning stability is a modulator of germline mutation rate variation across the human genome. Nat. Commun. 11, 1363 (2020).

15. Huang, Y. & Li, G.-M. DNA mismatch repair preferentially safeguards actively transcribed genes. DNA Repair 71, 82–86 (2018).

16. Li, F. et al. The histone mark H3K36me3 regulates human DNA mismatch repair through its interaction with MutSα. Cell 153, 590–600 (2013).

17. Huang, Y., Gu, L. & Li, G.-M. H3K36me3-mediated mismatch repair preferentially protects actively transcribed genes from mutation. J. Biol. Chem. 293, 7811–7823 (2018).

18. Wang, Y. et al. Histone H3 lysine 14 acetylation is required for activation of a DNA damage checkpoint in fission yeast. J. Biol. Chem. 287, 4386–4393 (2012).

19. Yazdi, P. G. et al. Increasing Nucleosome Occupancy Is Correlated with an Increasing Mutation Rate so Long as DNA Repair Machinery Is Intact. PLoS One 10, e0136574 (2015).

20. Salzberg, A. C. et al. Genome-wide mapping of histone H3K9me2 in acute myeloid leukemia reveals large chromosomal domains associated with massive gene silencing and sites of genome instability. PLoS One 12, e0173723 (2017).

21. Chong, S. Y. et al. H3K4 methylation at active genes mitigates transcription-replication conflicts during replication stress. Nat. Commun. 11, 809 (2020).

22. Belfield, E. J. et al. DNA mismatch repair preferentially protects genes from mutation. Genome Res. 28, 66–74 (2018).

23. Schuster-Böckler, B. & Lehner, B. Chromatin organization is a major influence on regional mutation rates in human cancer cells. Nature 488, 504–507 (2012).

24. Supek, F. & Lehner, B. Clustered Mutation Signatures Reveal that Error-Prone DNA Repair Targets Mutations to Active Genes. Cell 170, 534–547.e23 (2017).

25. Xia, B. et al. Widespread Transcriptional Scanning in the Testis Modulates Gene Evolution Rates. Cell 180, 248–262.e21 (2020).

26. Chen, X. et al. Nucleosomes suppress spontaneous mutations base-specifically in eukaryotes. Science 335, 1235–1238 (2012).

27. Supek, F. & Lehner, B. Differential DNA mismatch repair underlies mutation rate variation across the human genome. Nature 521, 81–84 (2015).

28. Exposito-Alonso, M. et al. The rate and potential relevance of new mutations in a colonizing plant lineage. PLoS Genet. 14, e1007155 (2018).

29. Frigola, J. et al. Reduced mutation rate in exons due to differential mismatch repair. Nat. Genet. 49, 1684–1692 (2017).

30. Heredia-Genestar, J. M., Marquès-Bonet, T., Juan, D. & Navarro, A. Extreme differences between human germline and tumor mutation densities are driven by ancestral human-specific deviations. Nat. Commun. 11, 2512 (2020).

31. Quadrana, L. et al. Transposition favors the generation of large effect mutations that may facilitate rapid adaption. Nat. Commun. 10, 3421 (2019).

32. Choi, J., Lyons, D. B., Kim, M. Y., Moore, J. D. & Zilberman, D. DNA Methylation and Histone H1 Jointly Repress Transposable Elements and Aberrant Intragenic Transcripts. Mol. Cell 77, 310–323.e7 (2020).

33. Lee, H., Popodi, E., Tang, H. & Foster, P. L. Rate and molecular spectrum of spontaneous mutations in the bacterium Escherichia coli as determined by whole-genome sequencing. Proc. Natl. Acad. Sci. U. S. A. 109, E2774–83 (2012).

34. Supek, F. & Lehner, B. Scales and mechanisms of somatic mutation rate variation across the human genome. DNA Repair 81, 102647 (2019).

35. Ossowski, S. et al. The rate and molecular spectrum of spontaneous mutations in Arabidopsis thaliana. Science 327, 92–94 (2010).

36. Wang, L. et al. The architecture of intra-organism mutation rate variation in plants. PLoS Biol. 17, e3000191 (2019).

37. Bobiwash, K., Schultz, S. T. & Schoen, D. J. Somatic deleterious mutation rate in a woody plant: estimation from phenotypic data. Heredity 111, 338–344 (2013).

38. Watson, J. M. et al. Germline replications and somatic mutation accumulation are independent of vegetative life span in Arabidopsis. Proc. Natl. Acad. Sci. U. S. A. 113, 12226–12231 (2016).

39. Arndt, P. F., Hwa, T. & Petrov, D. A. Substantial regional variation in substitution rates in the human genome: importance of GC content, gene density, and telomere-specific effects. J. Mol. Evol. 60, 748–763 (2005).

40. Duret, L. & Galtier, N. Biased gene conversion and the evolution of mammalian genomic landscapes. Annu. Rev. Genomics Hum. Genet. 10, 285–311 (2009).

41. Mugal, C. F. & Ellegren, H. Substitution rate variation at human CpG sites correlates with non-CpG divergence, methylation level and GC content. Genome Biol. 12, R58 (2011).

42. Youk, J., An, Y., Park, S., Lee, J.-K. & Ju, Y. S. The genome-wide landscape of C:G > T:A polymorphism at the CpG contexts in the human population. BMC Genomics 21, 270 (2020).

43. Long, H. et al. Evolutionary determinants of genome-wide nucleotide composition. Nat Ecol Evol 2, 237–240 (2018).

44. Fryxell, K. J. & Zuckerkandl, E. Cytosine deamination plays a primary role in the evolution of mammalian isochores. Mol. Biol. Evol. 17, 1371–1383 (2000).

45. Fryxell, K. J. & Moon, W.-J. CpG mutation rates in the human genome are highly dependent on local GC content. Mol. Biol. Evol. 22, 650–658 (2005).

46. Elango, N., Kim, S.-H., Vigoda, E. & Yi, S. V. Mutations of different molecular origins exhibit contrasting patterns of regional substitution rate variation. PLoS Comput. Biol. 4, e1000015 (2008).

47. Hodgkinson, A. & Eyre-Walker, A. The genomic distribution and local context of coincident SNPs in human and chimpanzee. Genome Biol. Evol. 2, 547–557 (2010).

48. Polak, P. et al. Cell-of-origin chromatin organization shapes the mutational landscape of cancer. Nature 518, 360–364 (2015).

49. Ha, K., Kim, H.-G. & Lee, H. Chromatin marks shape mutation landscape at early stage of cancer progression. NPJ Genom Med 2, 9 (2017).

50. Hung, S. et al. Mismatch repair-signature mutations activate gene enhancers across human colorectal cancer epigenomes. Elife 8, (2019).

51. Sabarinathan, R., Mularoni, L., Deu-Pons, J., Gonzalez-Perez, A. & López-Bigas, N. Nucleotide excision repair is impaired by binding of transcription factors to DNA. Nature 532, 264–267 (2016).

52. Chen, X. & Zhang, J. Yeast mutation accumulation experiment supports elevated mutation rates at highly transcribed sites. Proceedings of the National Academy of Sciences of the United States of America vol. 111 E4062 (2014).

53. Kovalchuk, I., Kovalchuk, O. & Hohn, B. Genome-wide variation of the somatic mutation frequency in transgenic plants. EMBO J. 19, 4431–4438 (2000).

54. 1001 Genomes Consortium. Electronic address: & 1001 Genomes Consortium. 1,135 Genomes Reveal the Global Pattern of Polymorphism in Arabidopsis thaliana. Cell 166, 481–491 (2016).

55. Carmel, L., Rogozin, I. B., Wolf, Y. I. & Koonin, E. V. Evolutionarily conserved genes preferentially accumulate introns. Genome Res. 17, 1045–1050 (2007).

56. Gorlova, O., Fedorov, A., Logothetis, C., Amos, C. & Gorlov, I. Genes with a large intronic burden show greater evolutionary conservation on the protein level. BMC Evol. Biol. 14, 50 (2014).

57. Mukherjee, D. et al. The role of introns in the conservation of the metabolic genes of Arabidopsis thaliana. Genomics 110, 310–317 (2018).

58. Liu, Y. et al. PCSD: a plant chromatin state database. Nucleic Acids Res. 46, D1157–D1167 (2018).

59. Kawakatsu, T. et al. Epigenomic Diversity in a Global Collection of Arabidopsis thaliana Accessions. Cell 166, 492–505 (2016).

60. McDonald, J. H. & Kreitman, M. Adaptive protein evolution at the Adh locus in Drosophila. Nature 351, 652–654 (1991).

61. Monroe, J. G. et al. Drought adaptation in Arabidopsis thaliana by extensive genetic loss-of-function. Elife 7, (2018).

62. Baggs, E. et al. Convergent Loss of an EDS1/PAD4 Signaling Pathway in Several Plant Lineages Reveals Co-evolved Components of Plant Immunity and Drought Response. Plant Cell (2020) doi:10.1105/tpc.19.00903.

63. Mi, H., Muruganujan, A., Ebert, D., Huang, X. & Thomas, P. D. PANTHER version 14: more genomes, a new PANTHER GO-slim and improvements in enrichment analysis tools. Nucleic Acids Res. 47, D419–D426 (2019).

